# The Effect of Previous Life Cycle Phase on the Growth Kinetics, Morphology and Antibiotic Resistance of *Salmonella* Typhimurium DT104 in Brain Heart Infusion and Ground Chicken Extract

**DOI:** 10.1101/326579

**Authors:** Jabari L. Hawkins, Joseph Uknalis, Tom P. Oscar, Jurgen G. Schwarz, Bob Vimini, Salina Parveen

## Abstract

Growth models are predominately used in the food industry to estimate the potential growth of select microorganisms under environmental conditions. The growth kinetics, cellular morphology and antibiotic resistance were studied throughout the life cycle of *Salmonella* Typhimurium. The effect of the previous life cycle phase (late log phase [LLP], early stationary phase [ESP], late stationary phase [LSP] and early death phase [EDP]) of *Salmonella* after reinoculation in brain heart infusion broth (BHI), ground chicken extract (GCE) and BHI at pH 5, 7 and 9 and salt concentrations 2, 3 and 4% was investigated. The growth media and previous life cycle phase had significant effects on the lag time (λ), specific growth rate (μ_max_) and maximum population density (Y_max_). At 2% and 4% salt concentration the LLP had the significantly (*P*<0.05) fastest μ_max_ (1.07 and 0.69 log CFU/mL/h, respectively). As the cells transitioned from the late log phase (LLP) to the early death phase (EDP), the λ significantly (*P*<0.05) increased. At pH 5 and 9 the EDP had a significantly (*P*<0.05) lower Y_max_ than the LLP, ESP and LSP. As the cells transitioned from a rod shape to a coccoid shape in the EDP, the cells were more susceptible to antibiotics. The cells regained their resistance as they transitioned back to a rod shape from the EDP to the log and stationary phase. Our results revealed that growth kinetics, cell’s length, shape and antibiotic resistance were significantly affected by the previous life cycle phase.

## Importance

*Salmonella* is a major foodborne pathogen and commonly associated with consuming undercooked poultry or products that have been cross contaminated with raw poultry. The disease caused by this bacterium is known as salmonellosis. The symptoms may include nausea, vomiting, abdominal cramps, diarrhea, fever, and headaches. Depending on the level ingested, strain characteristics, and host factors, the symptoms may last from 4 to 7 days. Adequate information is not available about its survivability under normal and unfavorable conditions. The present study provided valuable information about the growth and changes of *Salmonella* Typhimurium DT104 throughout its lifecycle under various antibiotic and environmental conditions (pH, salt concentration and ground chicken extract). The results of this study demonstrate that the previous life cycle should be considered when developing growth models of foodborne pathogens to better ensure the safety of poultry and poultry products

## Introduction

*Salmonella* is a Gram-negative, rod-shaped and non-spore forming bacterium. Salmonellosis, an infection caused by *Salmonella*, is one of the most common and widely encountered foodborne diseases, with tens of millions of human cases occurring worldwide annually (1). In the United States, there is an estimated one million salmonellosis cases annually causing 19,000 hospitalizations and 380 deaths (2). This has resulted in an increased interest to better understand the growth and changes of *Salmonella* under antibiotic and environmental stresses.

J. Wen et al. (3) suggested that current food safety studies that base the effectiveness of interventions on the ability to reduce foodborne pathogens in the stationary phase may be overestimated. They found that *Listeria monocytogenes* in the long term survival (LTS) phase has more resistance to thermal and high-pressure processing than in the stationary phase. As the availability of nutrients dissipated and *L. monocytogenes* transitioned into the LTS phase, the morphology changed from a rod to a coccoid shape. The coccoid shape increased its survivability and resistance to treatments. J. Wen et al. (4) also found that initial viable cell density (~10^6^ to ~10^10^ cfu/mL) and pH (5.36 – 6.85) of *L. monocytogenes* affected the transition to the LTS phase.

Although many models exist for the growth of bacteria in model food systems (5–12), there is limited information about the regrowth of bacteria at previous life cycle phases such as lag, log, stationary, death and long-term stationary (LTS) phase. The objective of this study was to investigate how the phase of *S.* Typhimurium DT104 at the log, stationary and death phase affects its transition back to the stationary phase. This study was conducted to observe the microbial behavior and the effect of previous life cycle phases at a constant temperature on the growth, cell morphology and antibiotic resistance patterns of *Salmonella* in inoculated brain heart infusion broth and ground chicken extract.

## Materials and methods

### Preparation of Bacterial Inoculum

A multiple-antibiotic resistant strain (ATCC 700408, American Type Culture Collection, Manassas, VA) of *Salmonella* Typhimurium definitive phage type 104 (DT104) was used in this study. This strain has an antibiotic resistance profile (chloramphenicol, ampicillin, tetracycline and streptomycin) that is found in chicken isolates (7) and has been used in similar growth modeling studies (6, 10). The stock culture of this strain was maintained at −80°C in brain heart infusion broth (BHI) (BBL, Difco, BD, Sparks, MD) that contained 15% (vol/vol) glycerol (Sigma, St. Louis, MO).

### Previous Life Cycle Phase

A growth curve was developed to identify the life cycle phases of DT104 (Fig. 1). One microliter of the stock culture was transferred into sterile BHI (5 mL) in a centrifuge tube (15 mL) followed by incubation at 37°C and 150 rpm from 0 to 770 h. This was carried out in three replicates and 50 μL of undiluted and appropriate 1:10 diluted (10^−1^ to 10^−5^) samples from cultures were spiral plated (Whitely Automated Spiral Plater, Microbiology International, Frederick, MD) onto tryptic soy agar (TSA) plates. The TSA plates were incubated at 37°C for 24 h and then enumerated. In Fig. 1, after the stationary phase, the culture had a slow and extended death phase. Since there is no time period of a stable density, it is referred to as the early death phase (EDP) instead of the LTS phase throughout the paper.

**Fig 1.**
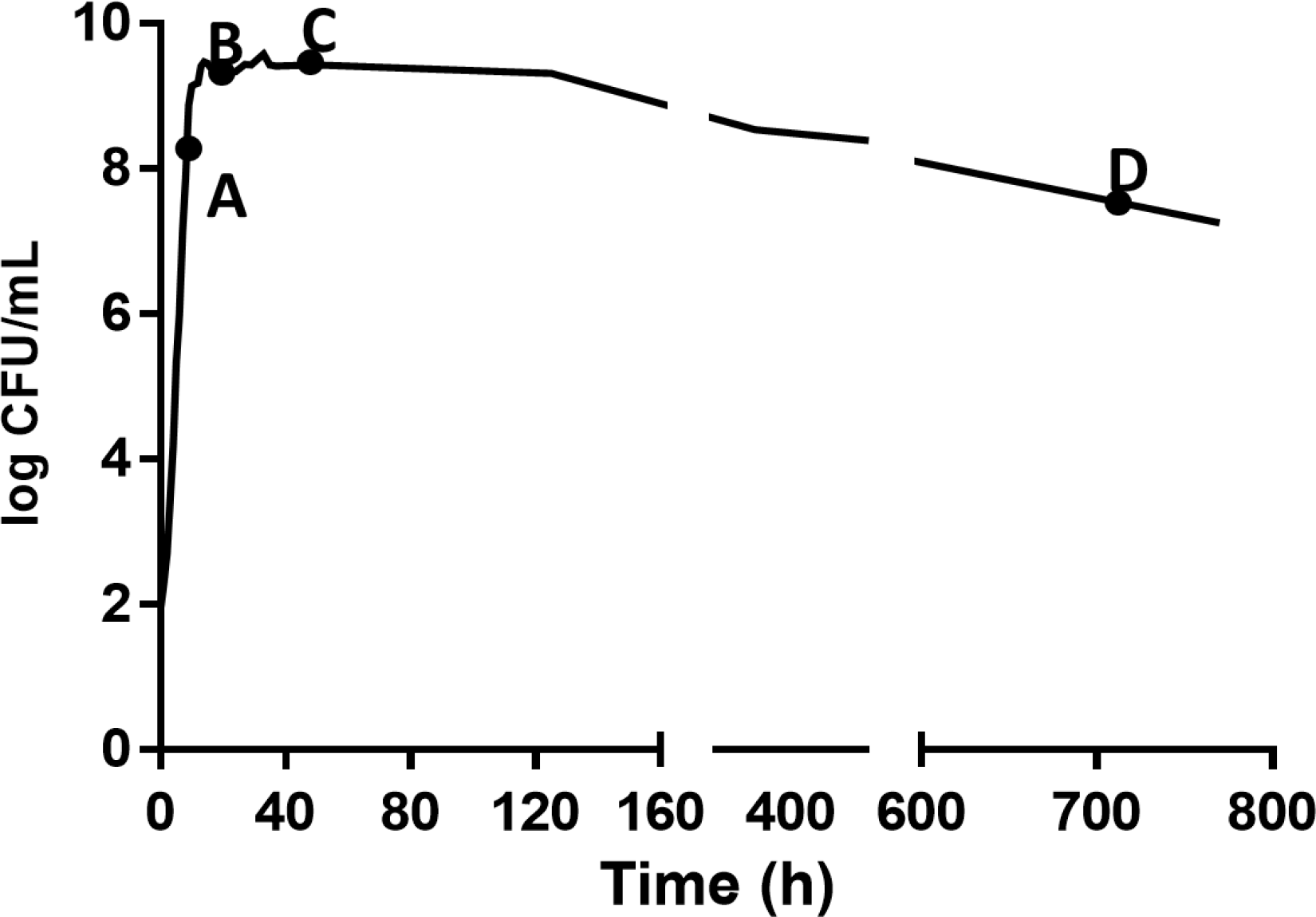
Growth of *S.* Typhimurium DT104 in brain heart infusion broth (BHI) at 37°C for different times to yield A (9 h, late log phase), B (24 h, early stationary phase), C (48 h, late log phase) and D (720 h, early death phase).

### Starter Culture

One microliter of the stock culture was transferred into sterile BHI (5 mL) in a centrifuge tube (15 mL) followed by incubation at 37°C and 150 rpm to obtain cells in the late log phase (LLP, 9 h), early stationary phase (ESP, 24 h), late stationary phase (LSP, 48 h) and early death phase (EDP, 720 h), respectively. The culture was centrifuged for 5 min at 5000 rpm and the pellet resuspended into 5 mL of the selected media as described below and adjusted to achieve a starting concentration of 10^4^ CFU/ml.

### Growth cultures

Analysis of cultures of DT104 for growth kinetic determinations were performed in centrifuge tubes (15 mL). Cultures grown under pH stress contained 5 mL of BHI adjusted to pH of 5, 7 or 9 with 1 N HCl or 1 N NaOH. Cultures grown under osmotic stress contained 5 mL of BHI adjusted to 2%, 3% and 4% NaCl (Difco Lab, MI, USA). DT104 was also grown in ground chicken extract (GCE) prepared as previously described in P. M. Fratamico et al. (13) with modifications. Fresh raw, 85% lean ground chicken was obtained from a local supermarket and high pressure processed (HPP) at 87,500 psi (603MPa) for 2.5 min. Sterile water (15 mL) was added to ground chicken (50 g) in a stomacher bag and pummeled for 1 min, the liquid extract was removed by centrifugation at 2100 × g for 5 min, and then filtered using a 0.22 μm filter, and frozen at −20°C until used. Growth cultures were incubated at 37°C and 150 rpm for 0 to 30 h.

### Microbial Analysis

At selected times post inoculation, depending on the age of the culture and growth culture medium of incubation, 50 μL of undiluted and appropriate 1:10 diluted (10^−1^ to 10^−5^) samples from growth cultures were spiral plated (Microbiology International, Frederick, MD) onto tryptic soy agar (TSA) plates. Sampling times for each growth condition (i.e. pH, salt concentration and GCE) were based on estimated lag time (λ), specific growth rate (μ_max_) and maximum population density (Y_max_) and were selected to produce a growth curve that accurately defined the lag, log and stationary phase over three log cycles of growth. The TSA plates were incubated at 37°C for 24 h.

### Predictive Growth Modeling

U.S. Department of Agriculture, Integrated Pathogen Modeling Program (IPMP) 2013 was used in this study. It is a free and easy-to-use data analysis platform for fitting primary models. Primary models include common growth and inactivation models, which can be used to analyze full growth curves, incomplete growth curves and inactivation/survival curves (14).

The data from the samples were assigned to IPMP 2013. The primary models evaluated were the full-growth Baranyi (15), reparametrized Gompertz (16), Huang (14) and Buchanan (17) three-phase linear. The Baranyi model, results not shown, was the most accurate and best fit for growth data and was used for modeling the data.

The Baranyi model is the following:

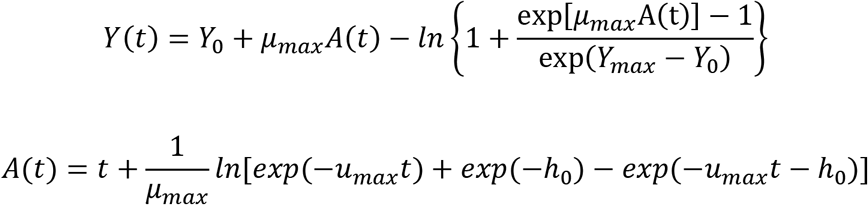

where y_o_, y_max_, y(t) are the bacterial population, in natural logarithm of bacteria counts, at initial, maximum and time t. μ_max_ is the specific growth rate; h_o_ is the physiological state of the microorganism under consideration.

#### 2.6 Scanning Electron Microscope (SEM) and Transmission Electron Microscope (TEM)

Scanning Electron Microscope (SEM) and Transmission Electron Microscope (TEM) was performed on cultures (20μL) of DT104 in different life cycle phases as described by J. Wen et al. (3). A one-way analysis of variance was used to determine the effect of the previous life cycle phase on the cell length of *S.* Typhimurium DT104. Pairwise comparisons were made by Tukey’s least significant difference test (α = 0.05) using Statistix 9 (Analytical Software, Tallahassee, FL).

### Antibiotic Susceptibility Disk Diffusion

Cultures of DT104 in different life cycle phases were tested for antibiotic susceptibility by the disc diffusion method on Mueller-Hinton agar (Sigma-Aldrich, Munich, Germany). The antibiotic sensitivity of the cultures of DT104 in different life cycle phases were evaluated according to classification guidelines suggested by the National Committee for Clinical Laboratory Standards (NCCLS). The cultures were tested for susceptibility to a panel of antibiotics including gentamicin (GM; 10 μg), sulphamethoxazole x trimethoprim (SXT; 25 μg), kanamycin (K; 30 μg), tetracycline (TE; 30 μg), nalidixic acid (NA; 30 μg), trimethoprim (TMP; 5 μg), ciprofloxacin (CIP; 5 μg), centriaxone (CRO; 30 μg), sulfisoxazole (G; 250 μg), chloramphenicol (C; 30 μg) and streptomycin (S; 10 μg) using the disk diffusion method (18,19). The zones of inhibition (ZOI) were measured and the isolates were classified as being susceptible, intermediate or resistant to the antibiotic.

### Statistical Analysis

Two-way analysis of variance (ANOVA) was used to determine the effect of the previous life cycle phase, growth medium (independent variables) and their interaction on the λ, μ_max_ and Y_max_ (dependent variables) of *S.* Typhimurium DT104 in various growth mediums (GCE and BHI) generated from the Baranyi model using GraphPad Prism 7 (GraphPad Software, San Diego, CA).

## Results

### Growth Models

The experimental data obtained at various previous life cycle phases and under environmental conditions and stresses was fitted to the full Baranyi growth model (Figures 2–4). The Baranyi model was a good fit to the growth data to accurately estimate the growth parameters (λ, μ_*max*_, and Y_*max*_) for statistical analysis. The lower and upper 95% confidence intervals were defined for individual predictions.

**Fig 2.**
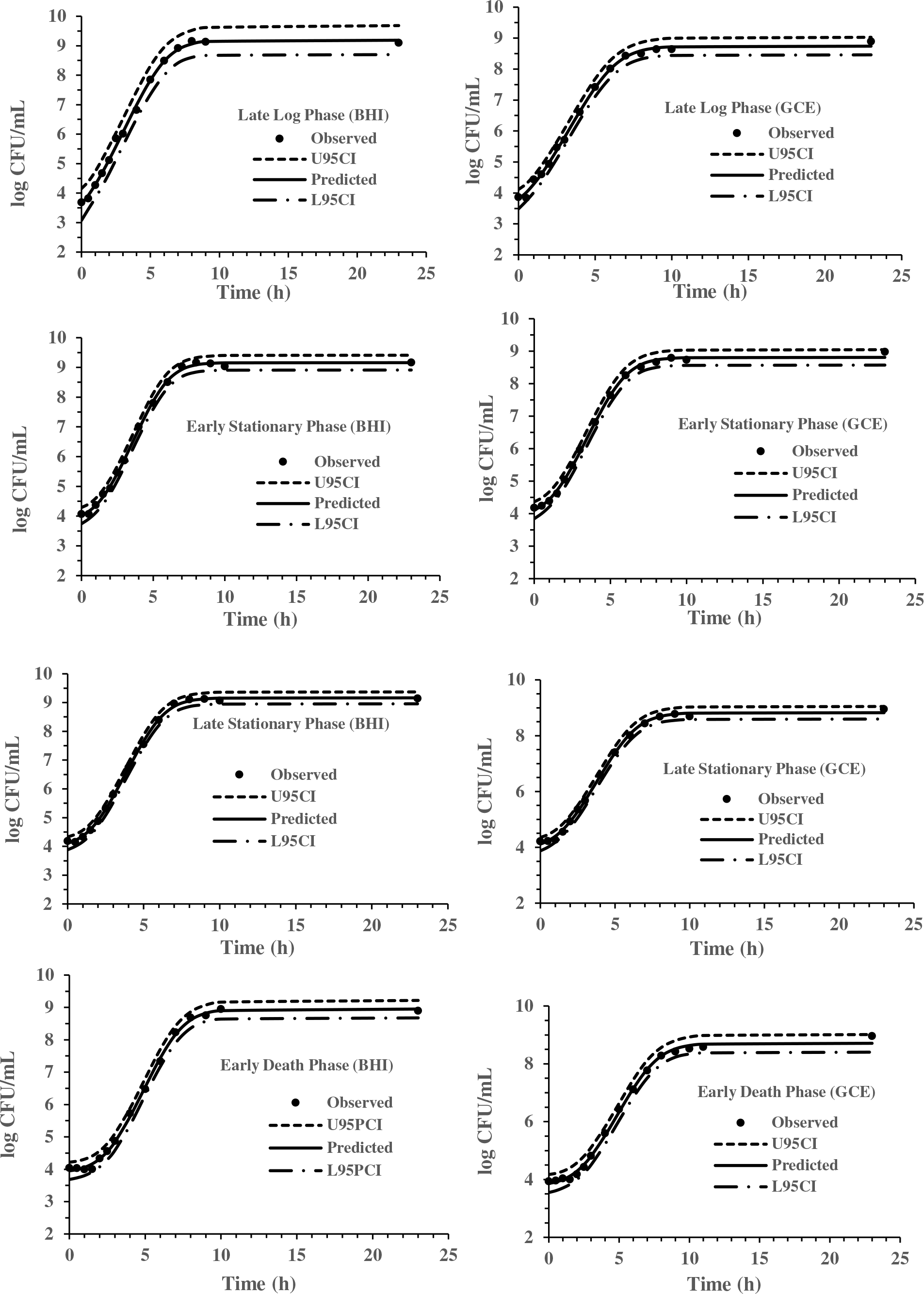
Baranyi Model for *S.* Typhimurium DT104 in brain heart infusion broth (BHI) and ground chicken extract (GCE) at 37°C. Observed, predicted, U95CI (upper 95% confidence interval) and L95CI (lower 95% confidence interval).

**Fig 3.**
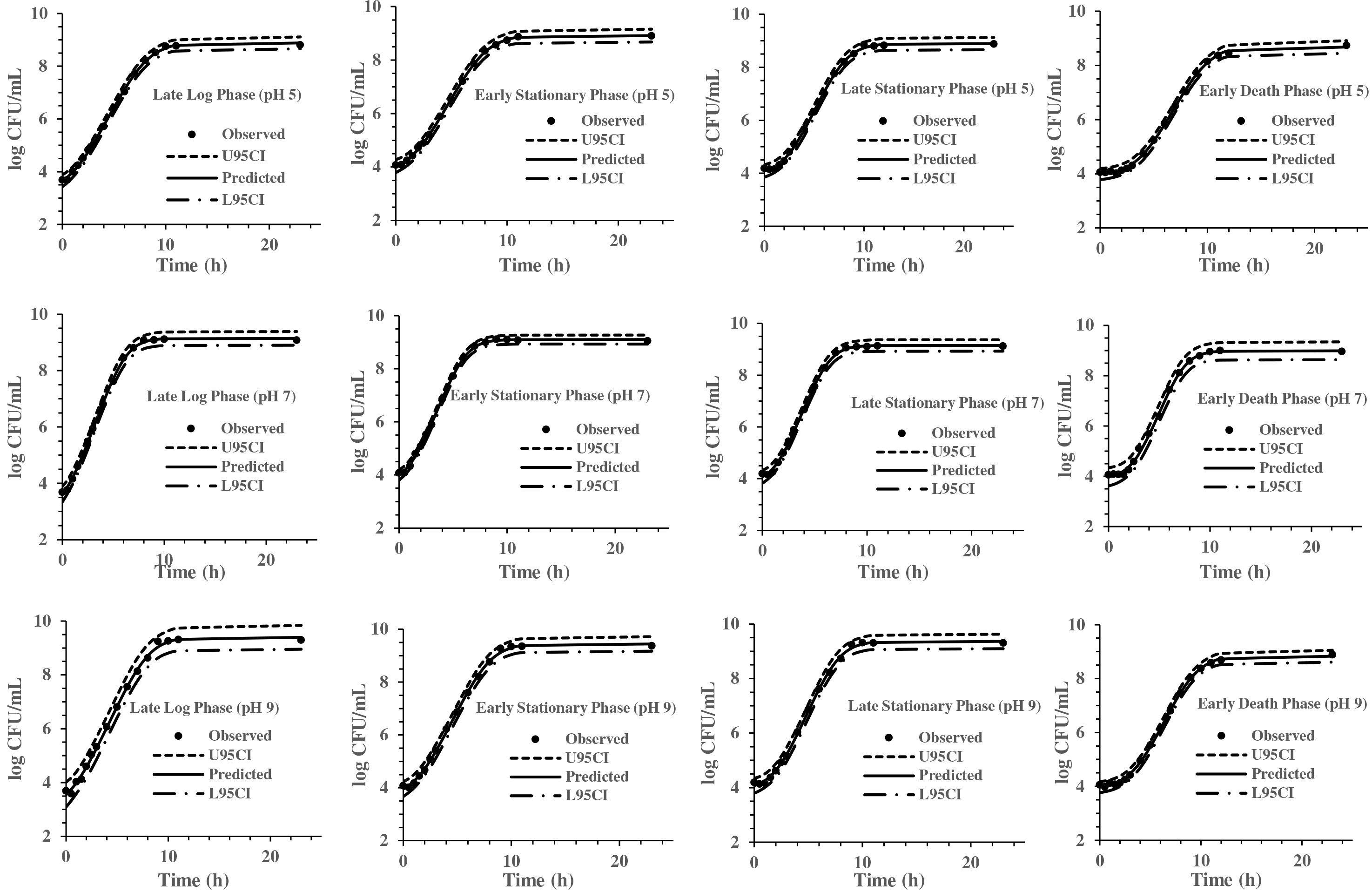
Baranyi Model for *S.* Typhimurium DT104 brain heart infusion broth (BHI) at pH 5, 7 and 9 at 37°C. Observed, predicted, U95CI (upper 95% confidence interval) and L95CI (lower 95% confidence interval).

**Fig 4.**
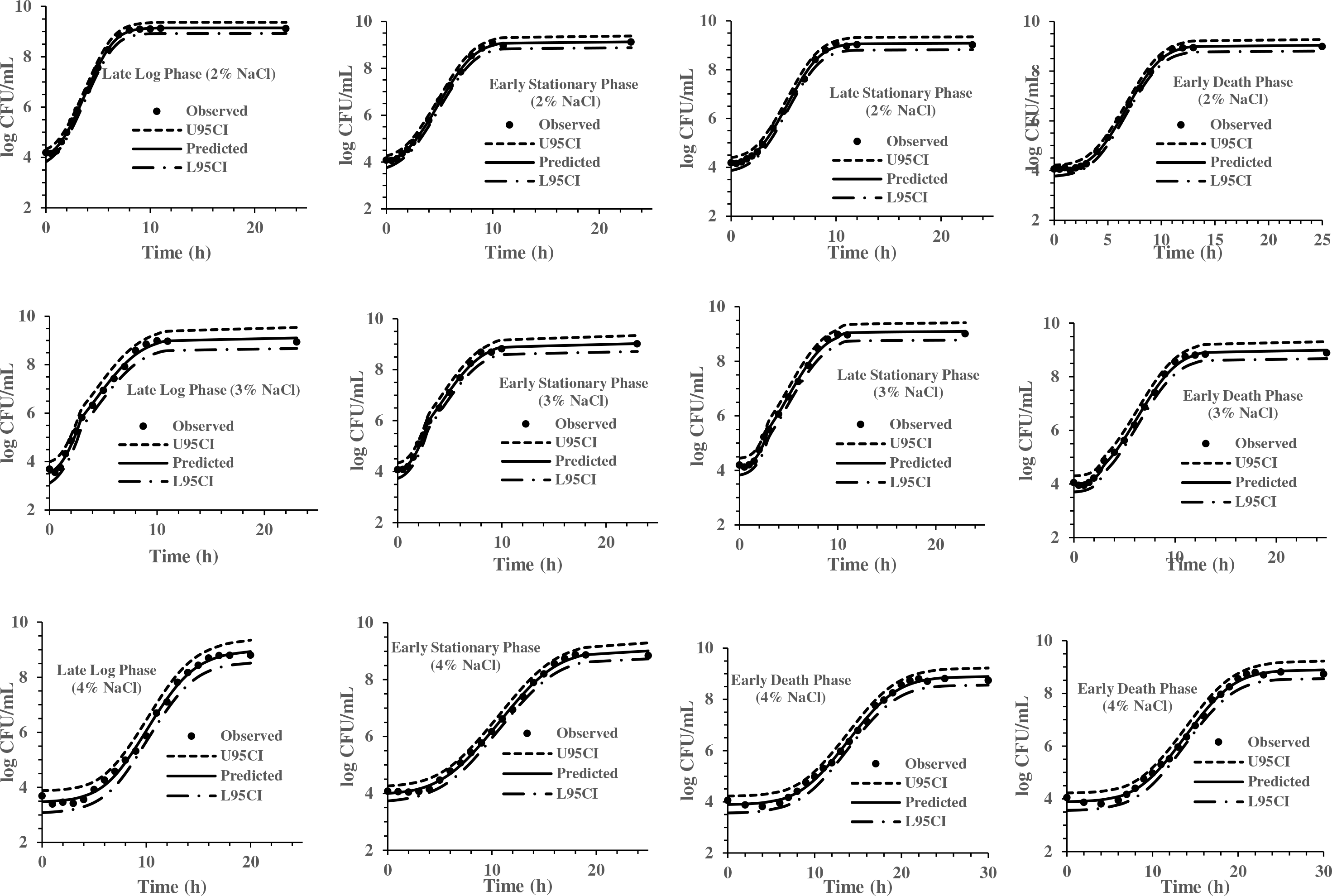
Baranyi Model for *S.* Typhimurium DT104 in brain heart infusion (BHI) at salt concentration 2, 3 and 4% at 37°C. Observed, predicted, U95CI (upper 95% confidence interval) and L95CI (lower 95% confidence interval).

### Brain Heart Infusion Broth vs Ground Chicken Extract

The cultures responded differently depending upon the phase of the culture. There was a significant (*P<0.05*) interaction between the medium and previous life cycle phase on the *Y*_*max*_. As seen in Fig. 5, the EDP in BHI had a lower (*P*<0.05) *Y*_*max*_ (8.95 log CFU/mL) in comparison to the LLP, ESP and LSP (9.19, 9.16 and 9.16 log CFU/mL, respectively). While, there was no significant (*P*>0.05) effect of the previous life cycle phases on the *Y*_*max*_ in GCE. There was no significant interaction between the media and previous life cycle phase on the λ and μ_*max*_ therefore only the main effects will be discussed. In BHI the λ significantly (*P* < 0.05) increased as the age of the previous life cycle phase increased except that the ESP λ equaled the LSP λ (LLP< ESP=LSP<EDP). There was also a significant (*P*<0.05) effect of the previous life cycle on the λ in GCE. The EDP had the significantly (*P*<0.05) longest λ at 2.47 h and there was a significant (*P*>0.05) difference between LLP and LSP of 0.19 h (Fig 5.). The LLP had a significantly (*P*<0.05) lower μ_*max*_ than ESP in both BHI (1.03 vs. 1.18 h^−1^, respectively) and GCE (0.96 vs. 1.10 h^−1^) (Fig. 5).

**Fig 5.**
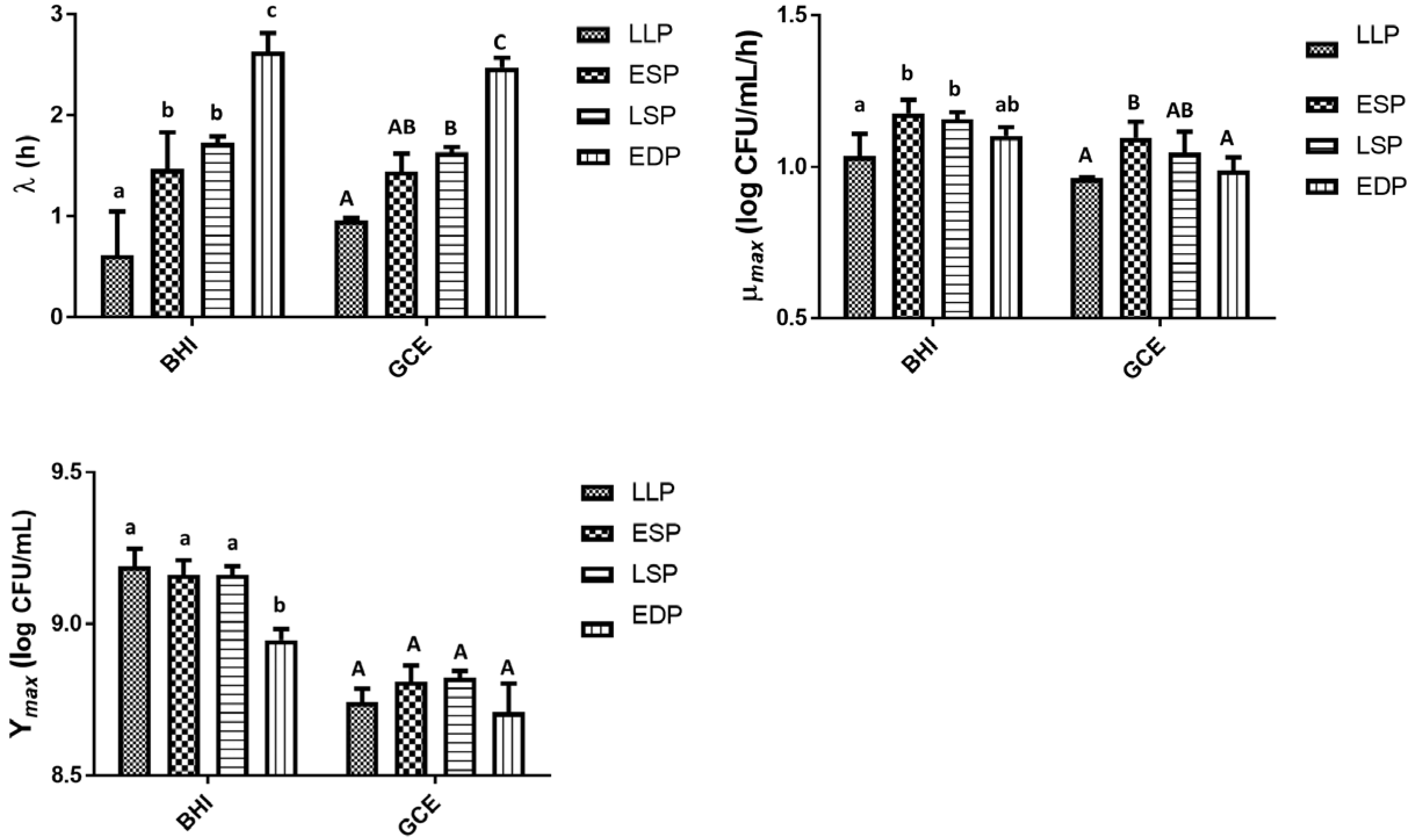
The lag time (λ), growth rate (μ_max_) and maximum growth (*Y*_*max*_) of *S.* Typhimurium DT104 as a function of the previous life cycle phase, LLP (late log phase), ESP (early stationary phase), LSP (late stationary phase) and EDP (early death phase) in brain heart infusion broth (BHI) and ground chicken extract (GCE). Different lowercase and uppercase letters within a cluster of bars differed significantly (*P < 0.05*).

### pH Levels

There was a significant (*P* < 0.05) interaction on the Y_*max*_ between the previous life cycle phase and various pH levels (Fig. 6). At pH 5 and 9 the EDP had a lower (*P*<0.05) Y_*max*_ than the LLP, ESP and LSP. While at pH 7 the previous life cycle phase had no significant (*P*>0.05) effect on the Y_*max*_. While there was a significant (*P* < 0.05) interaction on the Y_*max*_ there was none (*P* >0.05) on the λ and μ_*max*_. Similar to BHI and GCE, the λ significantly (*P* < 0.05) increased as the age of the previous life cycle phase (LLP, ESP, LSP and EDP) increased in both pH 5 (1.40, 1.90, 2.41 and 3.67 h, respectively) and pH 9 (1.04, 1.58, 2.13 and 3.56 h, respectively) (Fig. 6). At pH 7 the previous life cycle phase also had a significant (*P<0.05*) effect on the λ (LLP<ESP=LSP<EDP). The previous life cycle phase had no significant (*P>0.05*) effect on the μ_*max*_ regardless of the pH level (Fig. 6).

**Fig 6.**
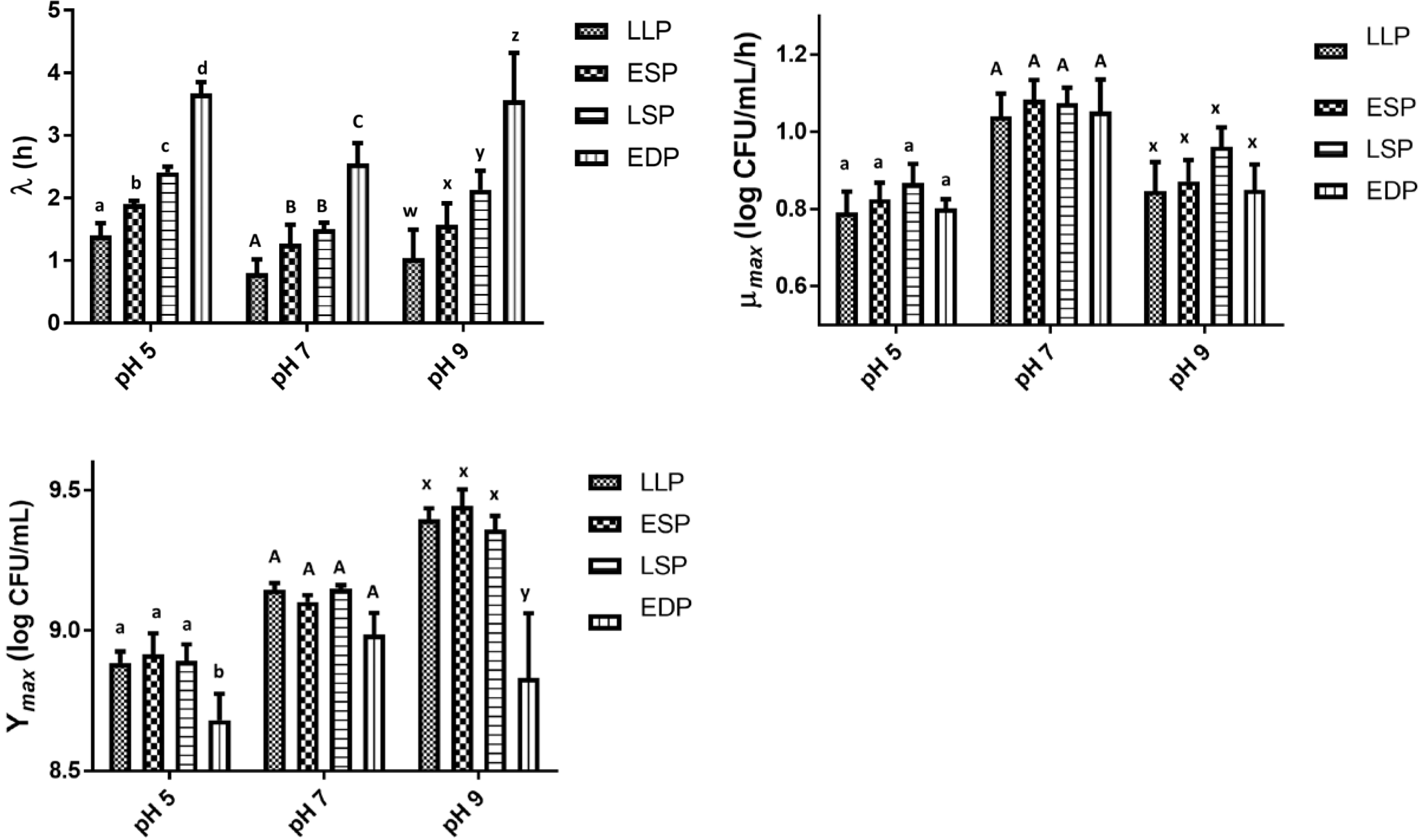
The lag time (λ), growth rate (μ_max_) and maximum growth (*Y*_*max*_) of *S.* Typhimurium DT104 as a function of the previous life cycle phase, LLP (late log phase), ESP (early stationary phase), LSP (late stationary phase) and EDP (early death phase) in brain heart infusion broth (BHI) at pH 5, 7 and 9. Different lowercase and uppercase letters within a cluster of bars differed significantly (*P < 0.05*).

### Salt Levels

In pertaining to the salt concentrations and the previous life cycle phase there was a significant (*P<0.05*) interaction on the μ_*max*_(Fig. 7). At 2% and 4% salt concentration, the LLP had the highest (*P*<0.05) μ_*max*_ and the previous life cycle phase had no significant (*P*<0.05) effect on the μ_*max*_ at 3% salt concentration. The previous life cycle phase and salt concentration had no significant (*P>0.05*) interaction on the λ and Y_*max*_. At 3% and 4% salt concentration the λ of LLP and ESP was not significantly (*P* >0.05) different and then the λ significantly (*P* <0.05) increased from the ESP to the EDP. However, at 2% salt the EDP had the longest (*P* > 0.05) λ (3.91 h). The salt concentrations had no significant (*P>0.05*) effect on the Y_*max*_ regardless of the previous life cycle phase (Fig. 7).

**Fig 7.**
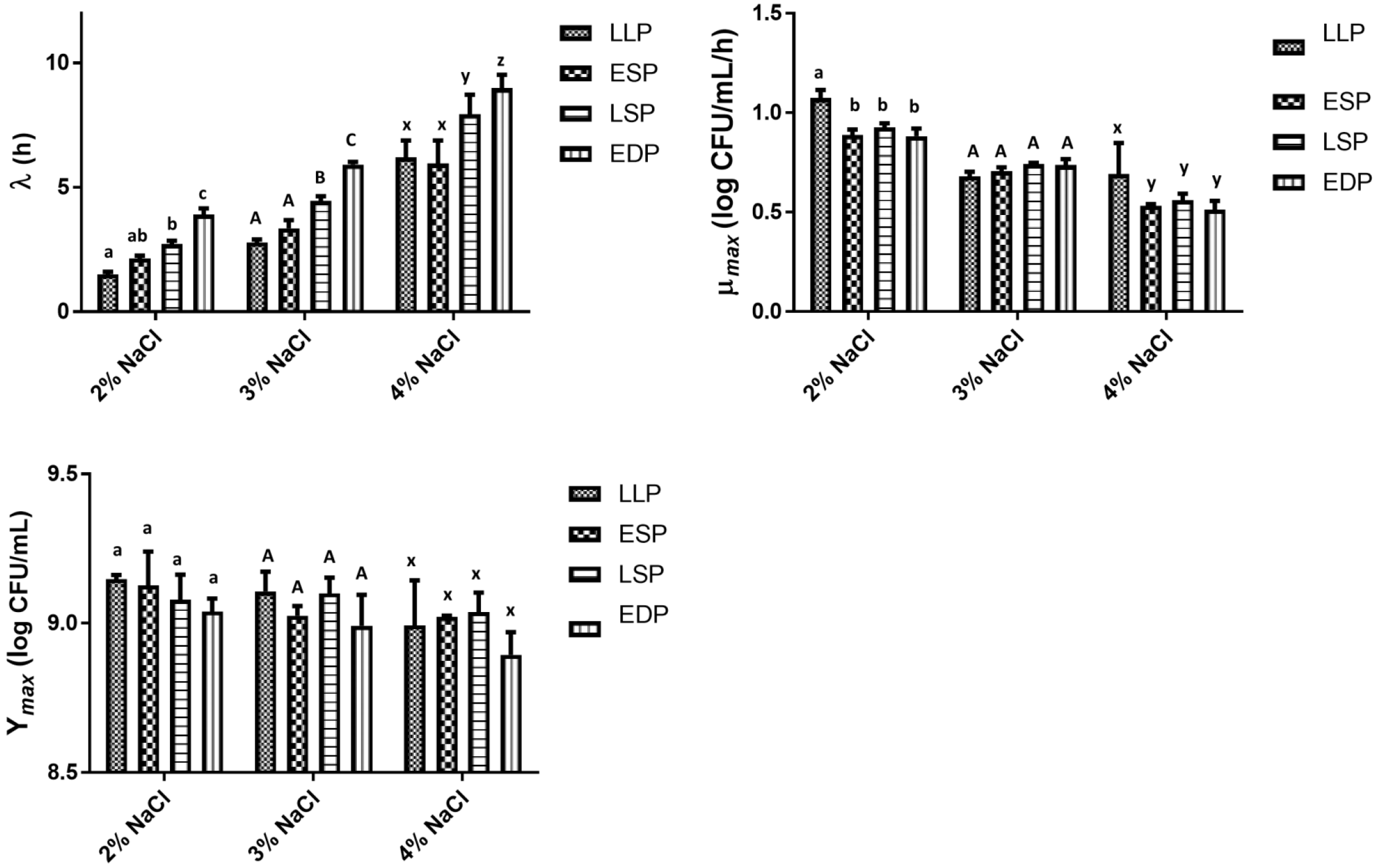
The lag time (λ), growth rate (μ_max_) and maximum growth (*Y*_*max*_) of *S.* Typhimurium DT104 as a function of the previous life cycle phase, LLP (late log phase), ESP (early stationary phase), LSP (late stationary phase) and EDP (early death phase) in brain heart infusion broth (BHI) at salt concentration 2, 3 and 4%. Different lowercase and uppercase letters within a cluster of bars differed significantly (*P < 0.05*).

### Scanning Electron Microscope and Transmission Electron Microscope

Representative photomicrographs of cells of DT104 in different phases are shown in Fig. 8. The SEM photomicrographs depicts the cells decreasing in size as they transitioned from the log phase (LP) to the EDP and size increased (recovered) transitioning back to the LP and stationary phase (SP) previously from the EDP. In the LP and SP the cells were rod-shaped while a cocci (spherical) shape appeared in the EDP. ANOVA of the SEM data revealed that cell length was significantly (*P<0.05*) affected by the growth phase with the LLP (2.79 μm) = LLP (previously EDP) (2.53 μm) > SP (previously EDP) (2.25 μm) = SP (2.14 μm) > EDP (1.81 μm).

**Fig 8.**
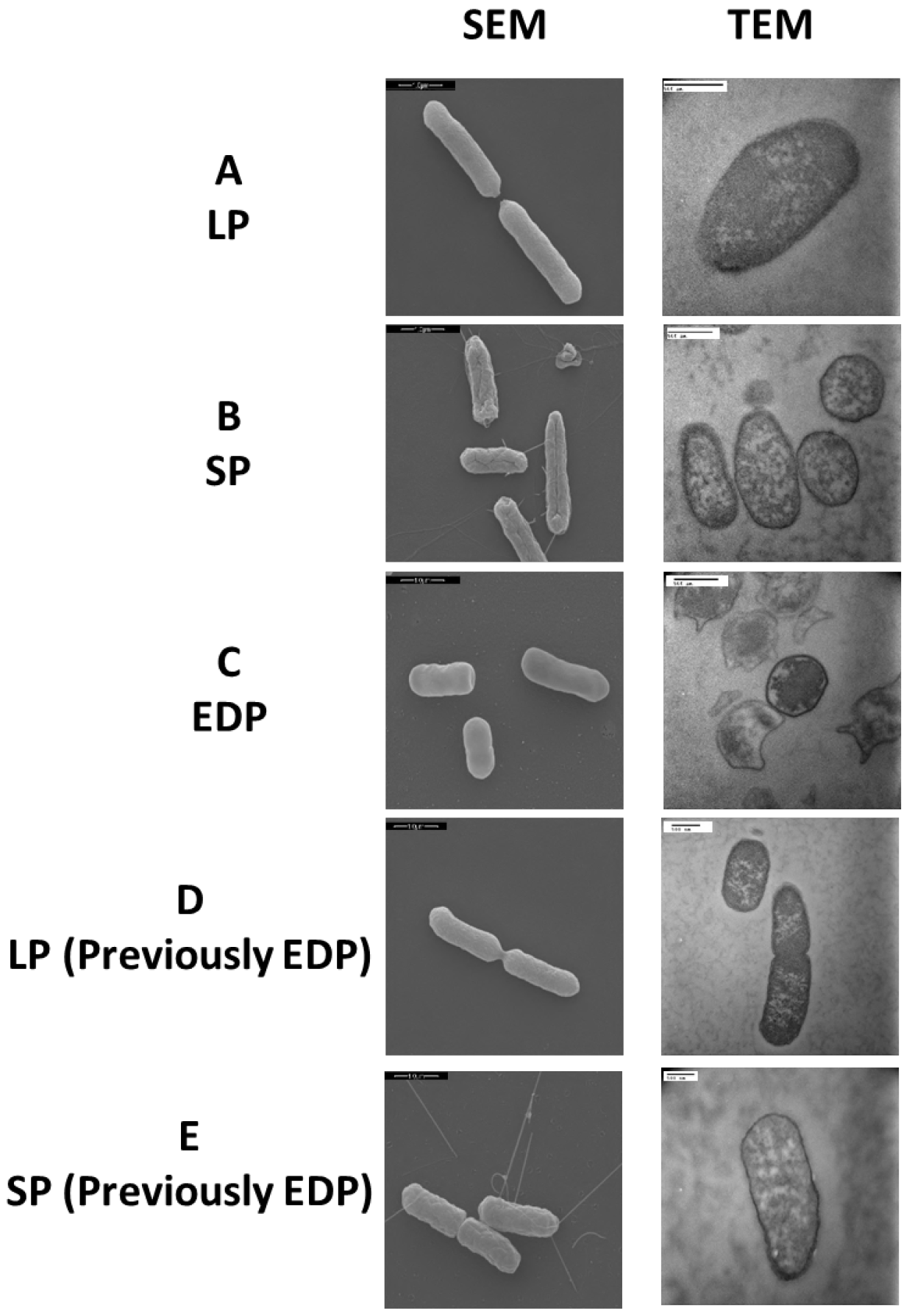
Scanning electron microscope (SEM) and transmission electron microscope (TEM) images of *S.* Typhimurium DT104 at different growth phases, LP (log phase), SP (stationary phase) and EDP (early death phase) in brain heart infusion broth (BHI) at 37°C.

### Antibiotic Susceptibility Disk Diffusion

Cells from the LP, SP and EDP grown in BHI were examined for their resistance to antibiotic treatment using susceptibility disk diffusion. DT104 from all phases were resistant to tetracycline (TE), sulfisoxazole (G), chloramphenicol (C) and streptomycin (S). In general, cells recovered from the EDP were more susceptible to the antimicrobial agents than the other phases especially for sulphamethoxazole × trimethoprim (SXT), in which the EDP was classified as susceptible to the antibiotic (ZOI 17.5 mm) but ESP, LSP and LLP (previously EDP) were intermediate. Cells from the LLP (previously EDP) are cells recovered from the EDP, regrown in fresh BHI to the LLP. Cells from the LLP (previously EDP) ZOI was similar to ESP and LSP when subjected to SXT, nalidixic acid (NA) and ciprofloxacin (CIP) except for gentamicin (GM), kanamycin (K) and ciprofloxacin (CIP) it was more susceptible and trimethoprim (TMP) and centriaxone (CRO) less susceptible. The LLP was more susceptible to GM, SXT, NA and CIP than ESP and LSP except for CRO, LLP was less susceptible.

## Discussion

### Growth Phases and Model Evaluation

It has been reported that the previous growth pH of a culture has a significant effect on the growth kinetics of *S.* Typhimurium (20) while the previous temperature and growth salt concentration is not a major factor affecting *S.* Typhimurium growth kinetics (8, 21). The objective of this study was to observe and analyze the significance of the previous life cycle phase on the growth kinetics of *Salmonella* Typhimurium. The model development phase of this study involved 32 growth curves conducted under combinations of pH, NaCl % and previous growth phase (LLP, ESP, LSP and EDP) identified in Fig. 1. IPMP 2013 was able to accurately fit the experimental data to the full Baranyi growth model. The smaller the value of the residual standard deviation, the better the fit of the regression curve to the data is (22). Growth at pH 5 (0.078) was the most accurately predicted data of the medium and the ESP (0.085) of the previous life phases.

Previous studies have demonstrated that after a short and rapid decline in the death phase, the culture maintains a stable density for an extended period of time (long term survival phase) (3, 4, 23). Exhibiting a different growth curve, in this study, the stationary phase (Fig. 1) lasted for a longer period of time in comparison of the stationary phase in J. Wen et al. (3). DT104 was not able to establish a LTS phase whereas J. M. N. Llorens et al. (23) and J. Wen et al. (3) maintained their own viable population at a constant population by continuous growth and death of cells. Although not establishing a LTS phase, the DT104 cells in the EDP died at a slow rate to survive for an extended period of time. When entering the death phase of the life cycle, the bacteria encounter an environment deprived of essential nutrients, i.e. carbon, phosphate and nitrogen. *Salmonella* may encounter periods of starvation in natural, host microenvironments and commercial environments. When deprived of carbon energy sources, *Salmonella* Typhimurium undergoes morphological changes as illustrated in Fig. 8 of the EDP which at times is referred to as starvation-stress response (SSR) (24).

To survive for an extended period of time, bacteria develop survival skills and strategies enabling them to persist in the environment until optimum growth conditions are met (25). Gram-negative bacteria such as *Vibrio cholerae, Escherichia coli* and *S.* Typhimurium may enter a state of dormancy or viable but not culturable (VBNC). In this state, cells are capable of metabolic activity but are unable to undergo cellular division to form a colony (24, 25). Additional research is needed to determine whether the cells in the EDP of Fig 1 are VBNC, which may explain the slow death rate. *S.* Typhimurium may also utilize nutrients that are not readily available in optimal growth conditions (24). In non-favorable growth conditions and environments, living cells utilize cryptic growth, which is the recycling of nutrients from dead cells for maintenance. In comparison to active and growing cells, cells transition from a rod shape to an efficient and smaller coccoid shape aiding in long term survival (3, 4, 25). During that transition, *S.* Typhimurium may use quorum sensing (QS) when responding to their own population density (26). When a high density cell culture senses limited nutrients, they undergo a form of programmed cell death, also known as bacterial apoptosis. The majority of the population enters a “death mode” while the surviving cells exit the death program after another signal is released perhaps from the lysed cells and may proceed to reproduce (27). This also might explain the survival of cells in the EDP. Quorum sensing is first initiated by small hormone-like molecules called autoinducers (AIs) that accumulate and monitor the environment (28). The LuxR homologue, SdiA achieves intercellular signaling and is capable of interspecies communication (29).

### Brain Heart Infusion and Ground Chicken Extract

BHI is a common nutrient rich broth used for the growth of *S.* Typhimurium and other fastidious and nonfastidious bacteria. The components (g/L) in BHI include: brain heart, infusion from solids (6), peptic digest of animal tissue (6), pancreatic digest of gelatin (14.5), dextrose (3), sodium chloride (5) and disodium phosphate (2.5). Chicken contains protein, including all of the essential amino acids, minerals (calcium, iron, magnesium, phosphorous, potassium, sodium, zinc and selenium), vitamins (niacin, pantothenic acid, vitamin B6 and B12, folate and folic acid (30). BHI contains 3.0 g/L of dextrose, whereas GCE does not and has a lower pH (6.4) than BHI (7.4). Overall, the GCE had a lower Y_max_ than BHI (8.77 vs 9.12 log CFU/mL, respectively). This may be due to the limited nutrients in the GCE, containing no carbohydrates, while BHI contains dextrose.

The present study and previous ones (6, 8, 10, 21) demonstrated that chicken based mediums can be used to develop models for the growth of *S.* Typhimurium in the place of traditional lab media. The λ of BHI after reinoculation of the ESP cells was 1.47 h which is similar to T. P. Oscar (5) at 1.25 h with similar growth conditions. T. P. Oscar (5) reported a slower μ_max_ at 0.86 log CFU/mL/h compared to 1.18 log CFU/mL/h in the present study. The λ of GCE after reinoculation of the ESP cells was 1.45 h while in T. P. Oscar (6), a study that developed predictive growth models for DT104 in ground chicken, had a λ of 1.39 and 1.66 h at 34° and 40°C, respectively. Using IPMP 2013 we determined the significance (*P*<0.05) of the previous life cycle phase and media on the growth kinetics of *S.* Typhimurium DT104.

A follow-up study will be performed on *S.* Typhimurium DT104 in GCE examining the global transcriptional profile of DT104 to provide insights on how this food matrix influences growth and survival and potentially identify specific genes that may be targeted for the development of control strategies. Previously, it has been difficult to perform transcriptomics in complex medium but due to technology advancement it has become possible instead of using a simple medium as BHI (13, 31). GCE may be a better growth medium to mimic a poultry environment in place of BHI for transcriptomic analysis of *Salmonella*.

### pH

*Salmonella* may encounter various pH levels in food production. Acid based antimicrobial solutions such as chlorine, organic acids, peroxyacetic and peracetic acid are commonly used in the poultry industry to reduce *Salmonella* and *Campylobacter* contamination on poultry and poultry products. Studies such as T. P. Oscar (20) and A. M. Gibson et al. (32) developed *Salmonella* growth models in broth media and demonstrated *Salmonella* ability to grow at low pH levels of 5.2 and 5.63, respectively. This study was carried out at a lower pH of 5 and demonstrated an ability of DT104 to grow in an unfavorable environment. Preliminary results (data not shown) demonstrated DT104 ability to grow at pH ≥ 4.5 but not at less than <4.5, this may be due to inability to maintain an adequate internal pH level. To the best our knowledge this is the first study that examined the effect of the previous life cycle and pH on the growth kinetics of a food pathogen. The starved cells in the EDP spent 2.27 h longer in the λ than LLP taking more time to adjust and adapt to pH 5 (Fig 3).

The growth kinetics have not frequently been studied at an alkaline pH. Alkaline cleaning agents are used in the food industry to remove biofilm, contamination, food, fat and protein from processing equipment (33, 34). Cleaners such as alkaline chlorinated cleaners work by temporarily imparting hydrophilic properties to stainless steel that decrease attachment to surfaces. While T. J. Humphrey (35) carried out a study demonstrating the inactivation of *Salmonella* at high pH and temperature, this is the first study to the best of our knowledge performing predictive growth models of *Salmonella* at high pH (pH 9).

### Salt

Salt is commonly added to food products to lower the water activity and inhibit microbial growth of pathogens and spoilage bacteria. In the presence of salt, the growth of bacteria is dependent upon the presence of osmoprotectants. In an abrupt shift of osmotic stress, cells accumulate harmful substrates such as K^+^ and glutamine. These substrates are replaced by proline, trehalose and glycine betaine, which are taken up from the medium or synthesized de novo by the cell to stimulate growth (36). Various studies (11, 37–39) have been carried out showing an extended λ of *Salmonella* as the salt concentration increases and the a_w_ decreases. The study by L. A. Mellefont et al. (38) carried out at a lower a_w_ demonstrated that the cells from the log phase after reinoculation had a longer relative lag time than the cells from the stationary phase in contrast to our study, where cells from the LLP after reinoculation had a shorter λ than the ESP but was carried out a higher a_w_ until at 4% NaCl cells from the LLP had a longer λ. These studies demonstrate that at concentrations higher than 4%, the cells from the LLP are not able to adjust to abrupt osmotic shifts. The cells with shorter λ may have better defense systems which prepares them to survive various environmental stresses without prior exposure. Further research is needed to better understand these mechanisms.

Similar to other studies (12), the μ_*max*_ decreased with the addition of salt (Fig. 7). This occurrence may be due to the initial response to osmotic shift with cells having 3 phases: dehydration, adjustment and rehydration. After respiration and ion transport are stopped from exposure to the shock, the respiration will resume and accumulate K^+^, glutamate and solutes. Unlike, K. Zhou et al. (36) there was an initial decrease in cell density in basic minimal medium at high salt concentration, DT104 did not have a decrease due to essential proteins and enzymes present in the BHI.

### SEM & TEM

Like *S.* Typhimurium DT104 in this study, other bacteria such as *L. monocytogenes* and *S.* Virchow have also shown a decrease in cell size (3, 40). DT104 transitioning back to the rod shape from the coccoid shape suggests that the cell formation is dependent upon the available nutrients. Starvation is known to induce a change in cell shape from rods to coccoid in *Arthrobacter crystallopoietes, A. globiformis, Rhizobium leguminosarum* and marine vibrio (25, 41–43). The EDP cells have a condensed cytoplasmic network in comparison to the other phases. This may explain the extended lag time of the subsequent EDP when introduced to a new environment. The coccoid shape may be formed by cell shrinkage and cytoplasmic condensation illustrated in Fig. 8 TEM. This may explain why the EDP cells require additional time to uptake available and essential nutrients and replenish fluidity in the cytoplasm.

### Antibiotic Susceptibility Disk Diffusion

To the best of our knowledge, this is the first study to compare the effect of life cycle phases on the antibiotic resistance profiles of viable cells of cultures after the stationary phase. The DT104 cells in the EDP exhibited less resistance to antibiotics than the cells from the LP and SP. The SP exhibited higher resistance to the antibiotics than the LP. A previous study by M. Tabak et al. (44) demonstrated *Salmonella* in a planktonic state was more resistant in the SP than the LP to triclosan. Although the aging cells in this study showed less resistance, a study by J. Wen et al. (3) demonstrated *Listeria monocytogenes* barotolerance and thermotolerance significantly (*P<0.001*) increased as the cells transitioned from the late log phase to the long term survival phase. However, a study by G. Juck et al. (45) demonstrated that the cells in the log phase exhibited higher resistance to HPP than cells in the stationary phase.

DT104 in the EDP phase may have exhibited less resistance to antibiotics due to the loss of plasmids after prolong incubation time and depletion of nutrients. Various studies have reported a modification of the plasmid pattern after long periods of incubation in marine water (46). It is possible that as the cultures continue to grow in the media the oxygen and nutrients become progressively more limited resulting in rendering the EDP cells more sensitive to antibiotics compared to early cells. *Salmonella* is capable of forming biofilms in environmental stresses for protection and to increase its long term survival. Biofilms, communities of bacterial cells, attach to one another and a surface. This structure may provide a higher degree of resistance to antibiotic agents, sanitizers and biocides. In the EDP, surviving in a state of low nutrients, DT104 formed visible clusters of biofilm along the wall and top layer of meniscus along the glass tube. However, the biofilm derived cells from the EDP were treated with antibiotics after disruption of the biofilm. This may have had an influence on the increased susceptibility to antibiotics compared to the other phases. After disruption, the potentially damaged cells from the EDP are exposed and vulnerable to the antibiotics.

## Conclusion

Using IPMP 2013, we accurately fitted the experimental data to the Baranyi full growth model to determine the significance of the previous life cycle phase (LLP, ESP, LSP and EDP) and media on the growth kinetics (λ, μ_*max*_, and Y_*max*_) of *S.* Typhimurium DT104. The results of this study showed that the previous life cycle phase has a significant effect on the λ, μ_*max*_, and Y_*max*_ and was also dependent upon the media. Overall, the X increased as the age of the previous life cycle phase increased. EDP may have spent a longer time in the lag phase repairing damaged cells, taking up available and essential nutrients and replenishing fluidity in the cytoplasm.

The results of this study suggest that not only the previous growth pH and temperature should be considered when developing growth models but also the previous life cycle phase. If the culture from the log phase has a shorter λ, studies that base their food growth model on foodborne pathogens from the stationary phase may be underestimated. Therefore, establishments that base their prerequisite programs or sanitation standard operating procedures on these models may be inadequate. Additional research on various serotypes of *Salmonella* is needed to prove this theory.

**Table 1.** Antibiotic disk diffusion of *Salmonella* Typhimurium DT104 as a function of previous life cycle phase

|  | Zone of Inhibition (mm) |  |  |  |  |
| --- | --- | --- | --- | --- | --- |
|  | LLP | ESP | LSP | EDP | LLP (Previously EDP) |
| **Aminoglycosides** |  |  |  |  |  |
| Gentamicin (10 µg) | S (20.0) | S (18.0) | S (18.0) | S (22.0) | S (21.0) |
| Kanamycin (30 µg) | S (17.5) | S (18.0) | S (17.5) | S (22.0) | S (19.0) |
| Streptomycin (10 µg) | R (--) | R (--) | R (--) | R (--) | R (--) |
| **Cephems** |  |  |  |  |  |
| Ceftriaxone (30 µg) | S (27.0) | S (28.0) | S (28.0) | S (31.0) | S (26.0) |
| **Folate Pathway Inhibitors** |  |  |  |  |  |
| Sulfisoxazole (250 µg) | R (--) | R (--) | R (--) | R (--) | R (--) |
| Sulphamethoxazole X Trimethoprim (25 µg) | S (16.5) | I (14.5) | I (14.0) | S (17.5) | I (14.0) |
| Trimethoprim (5 µg) | S (19.0) | S (18.0) | S (19.0) | S (22.5) | S (17.0) |
| **Phenicol** |  |  |  |  |  |
| Chloramphenicol (30 µg) | R (--) | R (--) | R (--) | R (--) | R (--) |
| **Quinolones and Fluoroquinolones** |  |  |  |  |  |
| Ciprofloxacin (5 µg) | S (32.5) | S (31.0) | S (32.0) | S (37.0) | S (33.0) |
| Nalidixic Acid (30 µg) | S (20.0) | S (19.5) | S (19.5) | S (22.0) | S (19.0) |
| **Tetracyclines** |  |  |  |  |  |
| Tetracycline (30 µg) | R (9.0) | R (9.5) | R (9.0) | R (9.5) | R (9.0) |

R = Resistant
I = Intermediate
S = Susceptible
LLP=late log phase; ESP= early stationary phase; LSP=late stationary phase; and EDP =early death phase

## Acknowledgments

This project was supported by the U.S. Department of Education, Title III, and Perdue Farms, Inc..

